# Self-Organized Amphiphiles are Poor Hydroxyl Radical Scavengers in Fast Photochemical Oxidation of Proteins Experiments

**DOI:** 10.1101/2020.10.19.345652

**Authors:** Zhi Cheng, Charles Mobley, Sandeep K. Misra, Joshua S. Sharp

**Affiliations:** Department of Biomolecular Sciences; Department of Chemistry and Biochemistry, University of Mississippi, Oxford, Mississippi 38677

## Abstract

The analysis of membrane protein topography using fast photochemical oxidation of protein (FPOP) has been reported in recent years, but still underrepresented in literature. Based on the hydroxyl radical reactivity of lipids and other amphiphiles, it is believed that the membrane environment acts as a hydroxyl radical scavenger decreasing effective hydroxyl radical doses and resulting in less observed oxidation of proteins. Here, we investigated the effect of bulk hydroxyl radical scavenging in FPOP using both isolated cellular membranes as well as detergent micelles. We found no significant change in radical scavenging activity upon the addition of disrupted cellular membranes with the membrane concentration in the range of 0-25600 cell/μL using an inline radical dosimeter. We confirmed the non-scavenging nature of the membrane with the FPOP results of a soluble model protein in the presence of cell membranes, which showed no significant difference in oxidation with or without membranes. The use of detergents revealed that, while soluble detergent below the critical micelle concentration acts as a potent hydroxyl radical scavenger as expected, additional detergent has little to no hydroxyl radical scavenging effect once the critical micelle concentration is reached. These results suggest that any scavenging effect of membranes or organized amphiphilic membrane mimetics in FPOP experiments are not due to bulk hydroxyl radical scavenging, but may be due to a localized scavenging phenomenon.

## INTRODUCTION

Membrane protein structure characterization is particularly challenging due to their amphiphilicity, insolubility in water, and the role of the membrane in protein structure. Mass spectrometry (MS) has recently become an increasingly attractive option for the structural characterization of membrane proteins due to its high sensitivity, low sample requirements, and its speed and ability to handle impure samples [1, 2]. Native MS has emerged as a complementary approach to X-ray crystallography and NMR in membrane protein studies through the use of membrane mimetics such as detergent micelles, bicelles, lipodiscs, liposomes, amphipols, and nanodiscs and soft ionization techniques [3, 4]. Even though native MS provides important insights into subunit stoichiometry, complex organization and interface stability, it also presents challenges due to ionization into and detection in the gas phase, which can lead to differences in the complexes detected [5]. In addition, different membrane mimetic systems can alter protein structure and/or suppress MS ionization [6]. MS-based methods for the structural characterization of proteins in their native membrane environment offer valuable insights.

Fast photochemical oxidation of proteins (FPOP), a method that generates hydroxyl radicals by UV laser-induced photolysis of hydrogen peroxide, has demonstrated the ability to react with a wide variety of amino acids on the surface of the proteins and label them without altering the native protein structure during labeling [7–9]. FPOP coupled with MS has been rapidly adopted to protein structural biology research in recent decades [7, 10–12]. However, FPOP coupled with MS for membrane protein analysis remains limited, in part due to the low extent of labeling that has been reported. One report suggested that only sulfur-containing amino acids, which are highly reactive, yield modified amino acids using FPOP [13]. Nanodiscs and detergents are commonly used to increase the hydroxyl radical labeling during the membrane protein FPOP studies [14, 15]. Yet, it was hypothesized that the membrane environment scavenged the hydroxyl radicals during FPOP, compromising the FPOP analysis.

We recently reported that it is possible to compensate for hydroxyl radical scavengers in solution by increasing the initial concentration of hydroxyl radicals generated [16]. However, it is currently unclear if the radical scavenging effect hypothesized to limit membrane protein oxidation is a general scavenging effect that can be compensated, a localized scavenging effect limited to the surface of the membrane or membrane mimetic, or if the limited reported membrane protein oxidation is due to another effect entirely. To evaluate the effect of membrane conditions on general hydroxyl radical scavenging in FPOP, we tested the general hydroxyl radical scavenging abilities of both isolated cell membranes, as well as Triton detergents with different critical micelle concentrations (CMCs) to determine the role of self-organization on membrane amphiphile hydroxyl radical scavenging.

## EXPERIMENTAL PROCEDURES

### Materials

SH-SY5Y cells were a generous gift from Dr. Jason Paris, University of Mississippi. Phosphate-buffered saline (PBS), sucrose, 4-(2-hydroxylethyl)-1-piperazineethanesulfonic acid (HEPES), potassium hydroxide (KOH), sodium carbonate (Na_2_CO_3_), myoglobin, [Glu]1-Fibrinopeptide B (GluB), 2-(N-morpholine)-ethanesulfonic acid (MES), catalase, methionine amide, Triton X-100, Triton X-405, calcium chloride (CaCl_2_) and formic acid were purchased from Sigma-Aldrich Corporation (St. Louis, MO, U.S.A.). Adenine, glutamine, dithiothreitol (DTT), LC/MS grade acetonitrile and water, anilinonaphthalene-8-sulfonic acid (ANS) were purchased from Fisher Scientific (Fair Lawn, NJ, U.S.A.). Hydrogen Peroxide (30%) was purchased from J.T. Baker (Phillipsburg, NJ, U.S.A.). Fused silica capillary was purchased from Digi-Key (MN, U.S.A.). Sequencing grade modified trypsin was purchased from Promega (Madison, WI, U.S.A.).

### Cell Membrane Extraction

The cells were washed three times with PBS then resuspended in a homogenization buffer containing 0.2 M sucrose and 20 mM HEPES, pH 7.5. Cell lysis was performed on ice using a probe sonicator on pulse mode for 25 seconds (5 seconds on and 10 seconds off intervals). Samples were then centrifuged for 10 min at 2000 ×g to remove the pellet containing the nuclear fraction and cellular debris. The supernatant was collected by ultracentrifugation at 135,000 ×g for 45 min at 4°C to remove cytoplasmic protein in the supernatant. The pellet was resuspended in 0.2 M Na_2_CO_3_, pH 11.0 and ultracentrifuged again to break up microsomes. Supernatant with excess cytosol was discarded and the pellet fraction resuspended in water and ultracentrifuged a third time to ensure all the additional protein was removed from the sample. Pellet fractions containing the cellular membrane were stored at −80°C [17].

### FPOP with Cell Membrane

FPOP solutions contained 50 mM sodium phosphate (pH 7.4), 2 mM adenine, 17 mM glutamine, 5 μM myoglobin, 5 μM GluB, various concentrations of MES (0, 5, 10, 20 mM), 100 mM hydrogen peroxide, and varying concentrations of isolated cell membranes (measured by initial cell count per μL). Samples were prepared in triplicate. 20 μl samples were pushed through a 100 μm inner diameter silica capillary and illuminated by a COMPex Pro 102 KrF excimer laser (Coherent Inc, CA) with energy setting of 120 mJ/pulse and fluence of 10.5 mJ/mm^2^ with the instrument layout as described in our previous publications [16, 18]. Adenine absorbance was measured in real time by an inline dosimeter (GenNext Technologies, CA) at 265 nm, with the change in adenine absorbance at 265 nm (ΔAbs_265_) being directly indicative of the effective radical concentration, and decreases in ΔAbs_265_ indicating increased hydroxyl radical scavenging [16]. Samples were collected into 25 μL of quenching solution containing 0.5 μg/μL of catalase and 0.5 μg/μL of methionine amide [19] .

### Membrane Mimetic - Detergent Sample Preparation

Four replicates of 33 μM of ANS dye and different concentrations of Triton X-405 and Triton X-100 (Figure S1) were mixed with the FPOP solution. ANS fluorescence intensity was measured by a plate reader (BMG Labtech, Germany) with excitation at 410 nm and for monitored at 490 nm for emission. CMC was determined to be the concentration at which the ANS fluorescence began to show a strong positive correlation with detergent concentration [20]. To determine the bulk scavenging capacity of detergent preparations, triplicate FPOP samples were prepared containing 50 mM sodium phosphate (pH 7.4), 2 mM adenine, 17 mM glutamine, 100 mM hydrogen peroxide, 5 μM GluB, and varying concentrations of X-405 (0, 0.3, 0.6, 0.9, 1.2, and 1.5 mM) or X-100 (0, 0.1, 0.2, 0.3, 0.4, 0.5 mM). Samples were processed for FPOP with laser energy of 110 mJ/pulse and the fluence was 11.2 mJ/mm^2^.

### LC-MS Analysis of samples

After FPOP of myoglobin, 50 mM of Tris containing 1 mM of CaCl_2_ (pH 8.0) and 5 mM dithiothreitol were added and heated at 95 °C for 15 min. Samples were immediately cooled on ice for 2 min. For the protein digestion, a 1:20 ratio of trypsin/protein was added into the samples and incubated at 37°C overnight with sample mixing. Trypsin was inactivated by heating samples at 95 °C for 10 min. Formic acid was added to 0.1% in the samples and were stored at −20 °C until LC-MS analysis. Tryptic-digested myoglobin was analyzed by an Orbitrap Fusion Tribrid mass spectrometer coupled to a Dionex Ultimate 3000 System (Thermo Fisher, CA). Samples were loaded onto a trap column first (Acclaim PepMap C18 5μm, 0.3mm × 5mm) and eluted onto a nano C18 column (Acclaim 3μm, 0.075 mm × 150mm) with at a flow rate of 0.3μL/min. Solvent A was 0.1% formic acid in water and solvent B was 0.1% formic acid in ACN. The gradient consists of 2% solvent B hold for 2 min, 2-35% solvent B over 30 min, 95% B over 2 min and held for 3 min, returned to 2% B over 1 min and held for 9 min. Byonic (v2.10.5, Protein Metrics, San Carlos, CA) was used to identify the peptides and determine the protein sequence coverage. Average oxidation per peptide were calculated with the help of Xcalibur, V.3.1 using the Equation 1, where *n*_*OX*_ denotes the average oxidation events per peptide and *I* represents the peak intensity of the peptides. The *n*_*OX*_ quantification for each peptide was calculated based on total unmodified and modified amino acid side chain as the dependent variable. In addition, the multiple oxidations per peptide was also taken into account for avoiding systematic underestimation of multiple oxidation events that can occur at higher radical doses [16, 21].

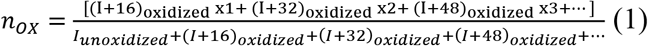

## RESULTS AND DISCUSSION

### Cellular Membranes as Hydroxyl Radical Scavengers

We added different concentrations of membrane extracted from SH-SY5Y cells into myoglobin FPOP samples (0, 400, 800, 1600, 3,200, 6,400, 12,800, and 25,600 cells/μL). Among all the measured cellular membrane concentrations, inline dosimetry data shows no significant scavenging effect caused by the cellular membrane. **(Figure 1A)**. FPOP of Glu-B was performed with or without membranes from 6400 cells/μL. MS data indicated no significant difference in scavenging activity and oxidation events of Glu-B peptide **(Figure 1B)**. We then performed FPOP of myoglobin in the presence or absence of membranes from 6400 cells/μL. Student’s *t*-test was used to compare the ΔA^265^ between the samples with or without membrane and the average oxidation event per peptide between the membrane and no membrane samples (*p* ≤ 0.05). As expected, there was no significant difference in the adenine absorbance change and the peptides from these two conditions did not show significant difference in oxidation **(Figure 1C)**. These results are in contrast to results obtained using increasing concentrations of MES, a soluble organic buffer that acts as a hydroxyl radical scavenger (**Figures S2 and S3, Supporting Information**). These results suggest that isolated cellular membranes do not act as bulk hydroxyl radical scavengers.

**Figure 1.**
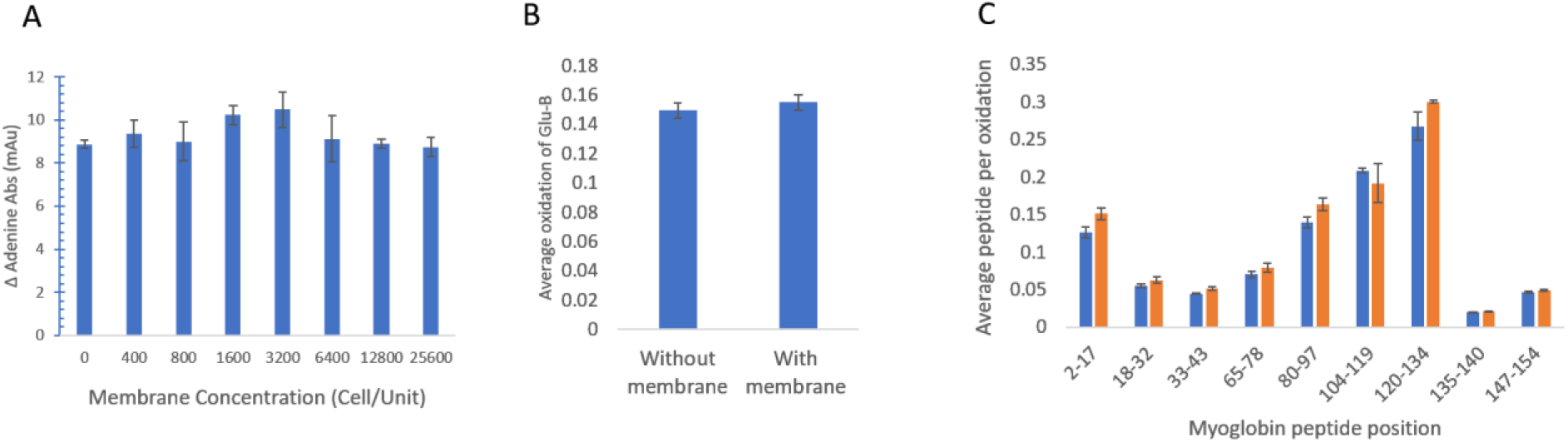
Cell membrane radical scavenging activity. **(A)** Different concentrations of isolated cellular membranes showed no significant changes in hydroxyl radical scavenging as monitored with in-line adenine dosimetry. **(B)** FPOP of the Glu-B peptide with or without 6400 cells/μL membrane showed no difference in its oxidation. **(C)** FPOP of myoglobin in the absence **(blue)** and the presence of 6400 cells/μL membrane **(orange)** showed no significant difference in peptide oxidation (α = 0.05).

### Effect of Self-Assembly of Amphiphiles on Radical Scavenging Capacity

Based on our results indicating that isolated cellular membranes were poor scavengers of diffusing hydroxyl radicals in bulk solution, we hypothesized that in the tightly packed cellular membrane environment, the hydroxyl radicals were not able to interact with the alkyl chains of the lipids that would have the strongest radical scavenging capacity; rather, the relatively inert head groups are all that are available to the diffusing radicals. We tested our hypothesis by observing the hydroxyl radical activity changes under different self-assembly conditions of non-ionic detergents. Triton-series detergents were used to create both soluble detergent solutions with no significant self-assembly, or conditions where added detergent would self-assemble into micelles, potentially protecting the more reactive phenyl group. The concentration above which added detergent self-assembles in water into micelles is known as the critical micelle concentration (CMC). By measuring the scavenging capacity of detergent concentrations above and below the CMC for each detergent, we could determine if the self-assembly of the amphiphiles impacts their ability to serve as hydroxyl radical scavengers, which could explain why organized cell membranes are poor bulk radical scavengers while individual lipids are effective scavengers [22].

Detergent CMC is reported within a narrow concentration range and fluctuates under different environmental conditions such as temperature, pressure, and the presence of electrolytes [23]. Therefore, we determined the approximate CMC of the detergents in our hands using an ANS dye assay. ANS fluoresces strongly at 490 nm when the dye is solubilized within the interior of a hydrophobic micelle [24]. Using ANS fluorescence, we determined that Triton X-405 has a CMC of ~0.9 mM, while Triton X-100 has a CMC of ~0.2 mM in our hands (**Figure 2)**. These CMC values are consistent with the reference ranges of CMCs as reported by the manufacturer (0.81 - 1.24 mM and 0.2 - 0.9 mM, respectively for Triton X-405 and Triton X-100). We then evaluated the hydroxyl radical scavenging effectiveness of different concentrations of detergent, both below and above the CMC. We used real-time adenine dosimetry to measure the bulk scavenging capacity of the detergent solution by monitoring changes in the ΔAbs_265_ (**Figure 2)**. When the detergent concentration was below its CMC, we found that increasing detergent concentration had a large effect on the ΔA265, indicating a substantial hydroxyl radical scavenging capacity. However, we found that this concentration-dependence of radical scavenging disappeared right around the CMC of the detergent (at ~0.62 mM for Triton X-405, and ~0.23 mM for Triton X-100). Above the CMC, no amount of increase in detergent concentration significantly decreased the ΔA265 of the adenine dosimeter, indicating that detergent organized into micelles did not act as an effective hydroxyl radical scavenger.

**Figure 2:**
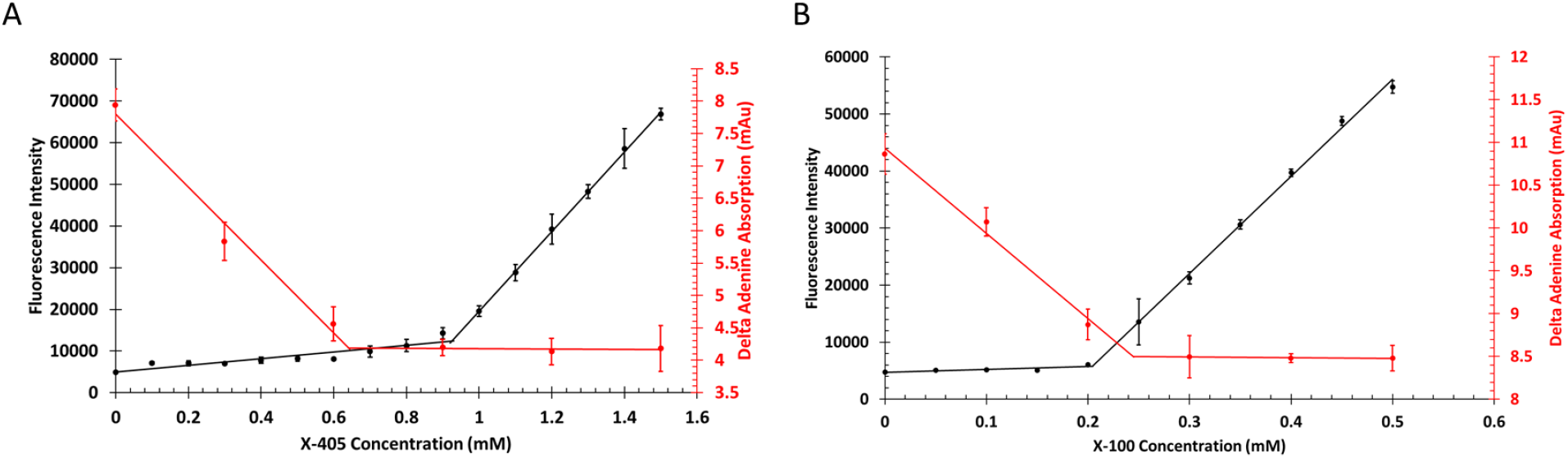
Hydroxyl radical scavenging activity is dependent on free detergent. The black line represents the fluorescence intensity of ANS dye. The red line represents hydroxyl radical scavenging activity by measuring ΔA_265_ of adenine with inline dosimeter. **(A)** Triton X-405 scavenges when detergent is free in solution, but once the CMC is reached additional detergent does not add to the scavenging capacity of the solution. **(B)** Triton X-100 shows a similar pattern, with the concentration region where additional detergent ceases to compete with adenine for the hydroxyl radical shifting to lower concentrations due to the lower CMC of Triton X-100.

## Conclusion

These findings indicate that the self-assembled structures of amphiphiles have a strong influence on their ability to act as scavengers of diffusing hydroxyl radicals. It has been widely reported that aromatic and aliphatic groups, such as, found in the hydrophobic groups of the Triton series of detergents, are very efficient hydroxyl radical scavengers [25]. Previous reports of FPOP of integral membrane proteins [13, 26] and the traditional concept of hydroxyl radical scavenging suggest that these amphiphiles act by reacting with hydroxyl radicals as they diffuse through solution reducing the effective concentration of hydroxyl radicals available to react with the protein target. However, the results presented here suggest that self-assembled membranes and membrane mimetics do not serve as efficient bulk hydroxyl radical scavengers. As shown in **Figure 1**, these membranes have absolutely no effect on the oxidation of proteins not directly associated with the membrane.

New ways of thinking about the role of the membrane environment in limiting the oxidation of integral membrane proteins are required. One possibility is that hydroxyl radical scavenging in general in FPOP needs to be reconsidered. In most FPOP experiments, the lifetime of the initial hydroxyl radical is very short (< 1 μs) [7, 8, 27] For integral membrane proteins that are spatially confined to the membrane environment, perhaps local scavenging effects dominate so that, even though the self-assembled membrane or membrane mimetic is a poor scavenger, the very high local concentration results in a high local scavenging capacity. However, other alternatives also exist. In cellular contexts, it is well-known that lipid oxidation products are common products of biological reactive oxygen species [28, 29] However, the role of lipid oxidation in FPOP of integral membrane proteins has not been characterized. Lipids may act as radical transfer partners for integral membrane proteins, where the initial oxidation occurs at a less reactive, solvent accessible amino acid and the radical is then transferred to a lipid (or lipid-soluble factor) where it could either be transferred back to a more reactive amino acid target (such as those reported to be oxidized in integral membrane proteins by FPOP) or terminated [30, 31] The oxidation of lipids by protein oxidation intermediates and products is a poorly-studied area [32] so it is unclear if this potential mechanism is significant.

## Acknowledgments

This work was supported by the National Institute of General Medical Sciences (R01GM127267). J.S.S. and S.K.M. acknowledge support for LC-MS analysis from the Glycoscience Center of Research Excellence (NIH P20GM103460).

## References

1. Khanal, A., et al., Pulsed hydrogen/deuterium exchange mass spectrometry for time-resolved membrane protein folding studies. J Mass Spectrom, 2012. 47(12): p. 1620–6.

2. Pan, Y., L. Brown, and L. Konermann, Hydrogen/deuterium exchange mass spectrometry and optical spectroscopy as complementary tools for studying the structure and dynamics of a membrane protein. International Journal of Mass Spectrometry, 2011. 302(1–3): p. 3–11.

3. Bolla, J.R., et al., Membrane Protein-Lipid Interactions Probed Using Mass Spectrometry. Annu Rev Biochem, 2019. 88: p. 85–111.

4. Johnson, D.T., L.H. Di Stefano, and L.M. Jones, Fast photochemical oxidation of proteins (FPOP): A powerful mass spectrometry-based structural proteomics tool. J Biol Chem, 2019. 294(32): p. 11969–11979.

5. Boeri Erba, E. and C. Petosa, The emerging role of native mass spectrometry in characterizing the structure and dynamics of macromolecular complexes. Protein Sci, 2015. 24(8): p. 1176–92.

6. Armstrong, K.L.R.a.D.W., Mechanism of signal suppression by anionic surfactants in capillary electrophoresis-electrospray ionization mass spectrometry. 1996.

7. Hambly, D.M. and M.L. Gross, Laser flash photolysis of hydrogen peroxide to oxidize protein solvent-accessible residues on the microsecond timescale. J Am Soc Mass Spectrom, 2005. 16(12): p. 2057–63.

8. Gau, B.C., J. Chen, and M.L. Gross, Fast photochemical oxidation of proteins for comparing solvent-accessibility changes accompanying protein folding: Data processing and application to barstar. Biochim Biophys Acta, 2013. 1834(6): p. 1230–8.

9. Xie, B., et al., Quantitative Protein Topography Measurements by High Resolution Hydroxyl Radical Protein Footprinting Enable Accurate Molecular Model Selection. Sci Rep, 2017. 7(1): p. 4552.

10. Zhang, B., et al., Implementing fast photochemical oxidation of proteins (FPOP) as a footprinting approach to solve diverse problems in structural biology. Methods, 2018. 144: p. 94–103.

11. Pan, Y. and L. Konermann, Membrane protein structural insights from chemical labeling and mass spectrometry. Analyst, 2010. 135(6): p. 1191–200.

12. Liu, X.R., M.M. Zhang, and M.L. Gross, Mass Spectrometry-Based Protein Footprinting for Higher-Order Structure Analysis: Fundamentals and Applications. Chem Rev, 2020. 120(10): p. 4355–4454.

13. Pan, Y., et al., Structural characterization of an integral membrane protein in its natural lipid environment by oxidative methionine labeling and mass spectrometry. Anal Chem, 2009. 81(1): p. 28–35.

14. Watkinson, T.G., et al., FPOP-LC-MS/MS Suggests Differences in Interaction Sites of Amphipols and Detergents with Outer Membrane Proteins. J Am Soc Mass Spectrom, 2017. 28(1): p. 50–55.

15. Lu, Y., et al., Fast Photochemical Oxidation of Proteins Maps the Topology of Intrinsic Membrane Proteins: Light-Harvesting Complex 2 in a Nanodisc. Anal Chem, 2016. 88(17): p. 8827–34.

16. Sharp, J.S., et al., Real Time Normalization of Fast Photochemical Oxidation of Proteins Experiments by Inline Adenine Radical Dosimetry. Anal Chem, 2018. 90(21): p. 12625–12630.

17. Xu, G., et al., Unveiling the metabolic fate of monosaccharides in cell membranes with glycomic and glycoproteomic analyses. Chemical Science, 2019. 10(29): p. 6992–7002.

18. Xie, B. and J.S. Sharp, Hydroxyl Radical Dosimetry for High Flux Hydroxyl Radical Protein Footprinting Applications Using a Simple Optical Detection Method. Anal Chem, 2015. 87(21): p. 10719–23.

19. Saladino, J., et al., Aliphatic peptidyl hydroperoxides as a source of secondary oxidation in hydroxyl radical protein footprinting. J Am Soc Mass Spectrom, 2009. 20(6): p. 1123–6.

20. Jumpertz, T., et al., High-throughput evaluation of the critical micelle concentration of detergents. Analytical biochemistry, 2011. 408(1): p. 64–70.

21. Kiselar, J. and M.R. Chance, High-resolution hydroxyl radical protein footprinting: Biophysics tool for drug discovery. Annual review of biophysics, 2018. 47: p. 315–333.

22. Buxton, G.V., et al., Critical-Review of Rate Constants for Reactions of Hydrated Electrons, Hydrogen-Atoms and Hydroxyl Radicals (.OH/.O-) in Aqueous-Solution. Journal of Physical and Chemical Reference Data, 1988. 17(2): p. 513–886.

23. Neugebauer, J.M., Detergents: an overview, in Methods in enzymology. 1990, Elsevier. p. 239–253.

24. Brito, R.M. and W.L. Vaz, Determination of the critical micelle concentration of surfactants using the fluorescent probe N-phenyl-1-naphthylamine. Analytical biochemistry, 1986. 152(2): p. 250–255.

25. Buxton, G.V., et al., Critical review of rate constants for reactions of hydrated electrons, hydrogen atoms and hydroxyl radicals (⋅ OH/⋅ O− in aqueous solution. Journal of physical and chemical reference data, 1988. 17(2): p. 513–886.

26. Li, K.S., L. Shi, and M.L.J.A.o.c.r. Gross, Mass Spectrometry-Based Fast Photochemical Oxidation of Proteins (FPOP) for Higher Order Structure Characterization. 2018. 51(3): p. 736–744.

27. Watson, I.J., ‡ Tiandi Zhuang,† Olga Charvátová,† Robert J. Woods,† and Joshua S. Sharp, Pulsed Electron Beam Water Radiolysis for Submicrosecond Hydroxyl Radical Protein Footprinting. Anal Chem, 2009.

28. Kanner, J., et al., Initiation of lipid peroxidation in biological systems. C R C Critical Reviews in Food Science and Nutrition, 1987. 25(4): p. 317–364.

29. Smith, W.L. and R.C. Murphy, Oxidized lipids formed non-enzymatically by reactive oxygen species. J Biol Chem, 2008. 283(23): p. 15513–4.

30. Janusz M. Gebicki, J.D., James Collins and Helen Tweeddale, Peroxidation of proteins and lipids in suspensions of liposomes, in blood serum, and in mouse myeloma cells. Acta Biochimica Polonica, 2000.

31. Gieseg, S., S. Duggan, and J.M. Gebicki, Peroxidation of proteins before lipids in U937 cells exposed to peroxyl radicals. Biochemical Journal, 2000. 350(1).

32. Davies, M.J., Protein oxidation and peroxidation. Biochem J, 2016. 473(7): p. 805–25.

