## Supplementary Information for "Self-Organized Amphiphiles are Poor Hydroxyl Radical Scavengers in Fast Photochemical Oxidation of Proteins Experiments"

### Supporting Information

X-405

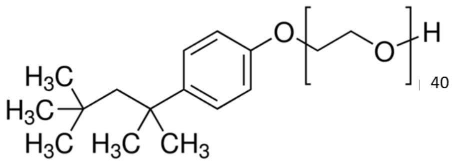

X-100

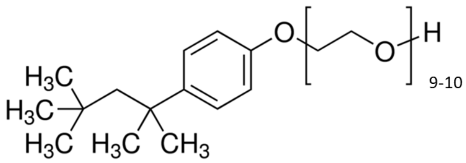

**Figure S1: Chemical structures of Triton X-405 and Triton X-100.** The detergents differ by the length of the ethylene oxide hydrophilic chain.

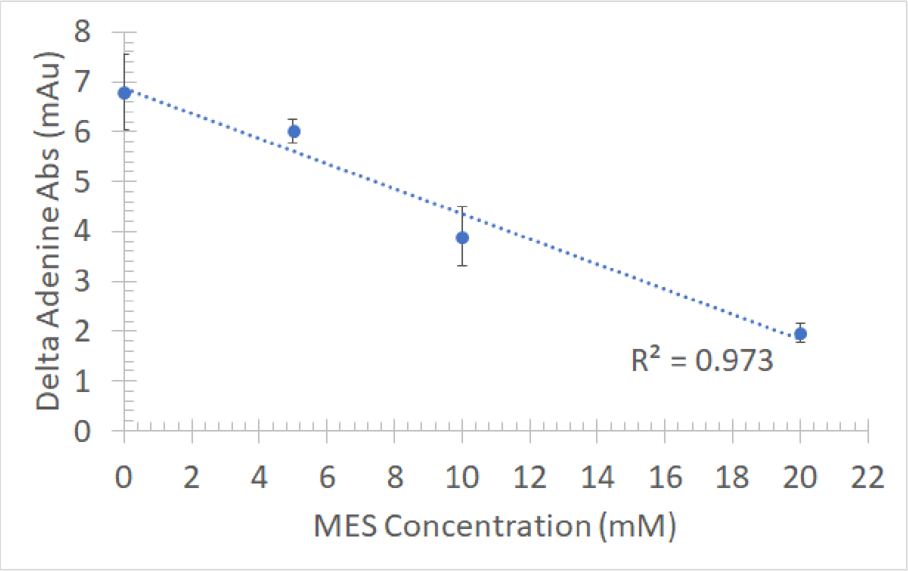


**Figure S2: Inline adenine dosimetry confirms radical scavenging of MES.** As the concentration of MES buffer increases, the $\Delta Abs_{265}$ (indicative of effective radical dose) decreases, showing MES buffer acted as a hydroxyl radical scavenger in these conditions.

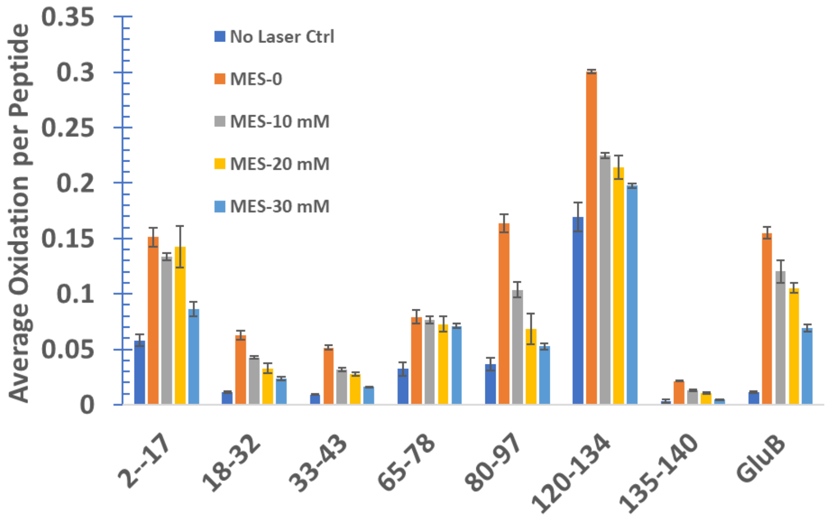


**Figure S3: Peptide and protein oxidation confirms radical scavenging of MES.** As the concentration of MES buffer increases, the amount of oxidation of all tryptic peptides from myoglobin and Glu-B peptide decreases, showing MES acted as a hydroxyl radical scavenger under these conditions.
